# Characterization of the Cell Division-Associated Peptidoglycan Amidase AmiA of *Chlamydia trachomatis*

**DOI:** 10.1101/2025.10.06.680639

**Authors:** Julia Dannenberg, Junghoon Lee, George W. Liechti, Christian Otten, Iris Löckener, Jula Reuter, Anna Klöckner, Sebastian Krannich, Tanja Schneider, Scot P. Ouellette, Beate Henrichfreise

## Abstract

*Chlamydia* is an obligate intracellular bacterium that differentiates between infectious, non-dividing EBs and non-infectious, dividing RBs. Pathogenic *Chlamydia* species are unusual in lacking a peptidoglycan sacculus, yet they do synthesize a transient and localized peptidoglycan structure at the divisome of the RB during their polarized division process. Although several studies have described the components of the chlamydial divisome necessary to generate peptidoglycan at a specific site on the membrane, less is understood about how the peptidoglycan structure is degraded to allow for the daughter cell to form and the division process to complete. Amidases are key components of the cell wall in model system bacteria as they catalyze the degradation and remodeling of peptidoglycan, including in the division septum. Here, we characterized the cell division-associated amidase, AmiA_Ct, of *Chlamydia trachomatis* both *in vitro* and *in vivo*. Our *in vitro* data show that AmiA_Ct is a *bona fide*, metal-dependent amidase capable of cleaving peptidoglycan. AmiA_Ct complemented an *E. coli* amidase deficient strain and supported the growth and separation of daughter cells. To assess the function of AmiA_Ct in *C. trachomatis*, we generated a transformant strain carrying an inducible CRISPR interference system targeting the *amiA* gene. Knocking down expression of *amiA* resulted in altered bacterial morphology, a reduction in infectious EBs, and the accumulation of peptidoglycan in the organisms. These data indicate a critical function for AmiA_Ct in the unique cell division process of *Chlamydia*.

**Importance:** Peptidoglycan is an important structural cell wall polymer that serves to give bacteria their shape and resistance to changes in extracellular solute concentrations. For *Chlamydia trachomatis*, an obligate intracellular pathogen that divides within a host cell, peptidoglycan is only used for cell division and is not a component of its cell wall. In this study, we characterize the function of a chlamydial amidase that helps degrade peptidoglycan during cell division. We show a critical function for amidase activity in facilitating changes to the peptidoglycan structure during chlamydial cell division that support normal growth and development of this pathogenic bacterium.

## Introduction

Bacteria of the genus *Chlamydia* are obligate intracellular pathogens of major public health concern globally, as they cause debilitating respiratory and sexually transmitted infections. During their developmental cycle, members of the Chlamydiae alternate between two morphologically distinct forms and depend on an eukaryotic host cell for proliferation [1]. As a consequence of reductive evolution and the relatively stable environment provided by the host cell, these microbes have significantly reduced their genome size and content [2, 3]. Whereas many chlamydial species have eliminated various nutrient biosynthesis pathways, pathogenic species (i.e., members of the Chlamydiaceae) have retained the genes necessary for peptidoglycan (PGN) biosynthesis even though they lack a PGN sacculus[4, 5]. PGN is an exoskeleton-like meshwork built from alternating sugar units that are crosslinked by peptide side chains, and, in most bacterial species, it encases the bacterial protoplast[6]. Biosynthesis spans three cellular compartments, starting in the cytoplasm with the synthesis of soluble precursors UDP-*N*-acetylglucosamine (UDP-Glc*N*Ac) and UDP-*N*-acetylmuramyl (UDP-Mur*N*Ac)-pentapeptide [7]. At the inner leaflet of the cytoplasmic membrane, UDP-GlcNAc is linked to UDP-MurNAc bound to the membrane carrier bactoprenol to form the precursor molecule lipid II [8]. Lipid II is flipped across the cytosolic membrane and integrated into the preexisting PGN meshwork by transpeptidation and transglycosylation carried out by penicillin binding proteins (PBPs) and SEDS proteins, respectively [9–11].

The fact that *Chlamydia* species synthesize PGN without an encasing sacculus raises the question why the intracellular organism has kept an energy-intense biosynthetic pathway for a seemingly unneeded molecule. However, small amounts of PGN could be detected in *Chlamydia* [12, 13]. Compared to free-living bacteria, chlamydial species employ alternative strategies to synthesize essential constituents of PGN, e.g., the biosynthesis of D-amino acids and *meso*-diaminopimelic acid (*m*DAP) [14, 15]. *Chlamydia* spp. have retained homologs of monofunctional transpeptidases PBP2 and PBP3 that are likely to incorporate PGN precursor lipid II into the already existing PGN meshwork [16, 17] and monofunctional carboxypeptidase PBP6 [16–18]. Transglycosylation is likely achieved by the SEDS proteins FtsW and RodA [19, 20].

For free-living bacteria, PGN represents the major osmoprotective and shape-determining element, and septal PGN plays a central role in cell division processes. In the model organism *E. coli*, PGN is constantly subjected to synthesis, remodeling, and degradation processes during cell growth and division. Upon initiation of cell division, a shared PGN septum is formed between nascent daughter cells, which is hydrolyzed during cell separation by cell division amidases [21, 22]. Even though members of the Chlamydiaceae do not synthesize a classical cell wall, PGN is needed for chlamydial cell division. The process is initiated by a polarized budding mechanism [23–25]. At the interface of the growing bud and the mother cell, a PGN disc is synthesized and subsequently altered into a ring structure [12, 13]. As the septum between dividing cells is constricted, the PGN ring is subject to constant remodeling reducing its diameter and is degraded at the end of the process.

This transient PGN is essential for chlamydial cell division, and the maintenance of the interconnected processes of PGN synthesis, remodeling, and recycling is absolutely required for proliferation. While novel PGN precursors are incorporated into the existing meshwork by transglycosylation, PGN is likely crosslinked by homologs of monofunctional transpeptidases PBP2 and PBP3 and subject to regulation by a homolog of monofunctional carboxypeptidase PBP6 [18]. Besides biosynthesis, PGN remodeling also requires catabolic processes for the PGN ring to expand and reduce its diameter. As a result of the genetic reduction that has streamlined their genomes to adapt to the intracellular niche, chlamydial species have developed alternative strategies to degrade PGN compared to those utilized by free-living bacteria. Cleavage of the glycan backbone in PGN is achieved by lytic transglycosylases SltY, MltA, MltB and others in *E. coli* [26–28]. In *Chlamydia*, no homologs of these enzymes exist, although a SpoIID homolog was shown to have lytic transglycosylase activity in *Chlamydia*-related bacteria [29].

In *E. coli* cell division, amide bonds of peptides crosslinking the glycan backbone are hydrolyzed by cell division amidases. The Gram-negative model organism encodes three zinc-dependent periplasmic amidases AmiA, AmiB, and AmiC, and their activity is tightly regulated requiring activation by LytM-domain containing proteins [30]. Until activation, the active sites of *E. coli* (Ec) AmiA, AmiB, and AmiC are occluded by an inhibitory helix structure. In AmiA_Ec, through interaction of protein activators with an interaction helix, a conformational change is forced upon the inhibitory helix occluding the active center, resulting in its release and in the exposure of the catalytic site. This regulatory process ensures that PGN degradation only occurs in a tightly controlled manner without compromising PGN and cell integrity. In members of the Chlamydiae, AmiA has been retained as a single cell division amidase homolog. AmiA from *Chlamydia* (*C.) pneumoniae* (AmiA_Cp) is functionally conserved and represents a novel penicillin target with dual activity as cell division amidase and carboxypeptidase [31]. In contrast to *E. coli* AmiA, the chlamydial ortholog degrades precursor molecule lipid II in addition to PGN *in vitro* [31]. Curiously, no mechanisms regulating chlamydial amidase activity are known, as homologs of LytM-domain containing proteins are not encoded in chlamydial species. This raises the question of how these microbes maintain PGN integrity during sensitive PGN remodeling processes that are tightly interlinked with cell division. Due to the unique bifunctional activity of AmiA_Cp and its role in catabolic PGN processes, we presume the enzyme to have a central function in sustaining a continuous cycle of PGN biosynthesis, remodeling, and recycling [31].

Most likely, PGN recycling is favorable to reduce the metabolic burden of *de novo* PGN biosynthesis: reusing PGN fragments is an energy-efficient strategy. In *E. coli*, PGN material is processed by a large multi-enzyme complex, and degradation products enter a new biosynthetic process. The PGN recycling machinery in genetically condensed *Chlamydia* spp. is significantly reduced but involves a homolog of multi-enzyme transporter OppABCDF in *C. trachomatis*, which transports PGN-derived peptides into the cytoplasm [32]. Recently, we characterized the *C. trachomatis* NlpC/P60-domain containing protein YkfC and demonstrated its importance for PGN recycling. By acting as an endopeptidase on the L-Ala-D-Glu-mDAP tripeptide, cleaving the non-canonical γ-D-Glu-mDAP bond exclusive to PGN, the enzyme facilitates reentry of PGN components into a new biosynthetic cycle [33]. Since PGN fragments are recognized as pathogen-associated molecular patterns (PAMPs) activating the host’s innate immune system, *Chlamydia* spp. are constantly at risk of being detected and eliminated. Specifically, the PGN-derived tripeptide L-Ala-D-Glu-mDAP represents the minimal motif of host innate immune factor NOD1 [34]. Thus, recycling of PGN fragments avoids shedding of immunogenic material into the surroundings and enables the pathogen to minimize its immunostimulatory profile [33].

In this work, we aimed to broaden our understanding of chlamydial cell division and the roles that the PGN ring and cell division amidase AmiA play in this highly coordinated process. We identified and characterized the AmiA homolog in *C. trachomatis* (AmiA_Ct), and our work revealed remarkable differences in the active site architecture between AmiA_Ct and the previously characterized AmiA_Cp [29]. Moreover, we show AmiA_Ct to be a monofunctional amidase in contrast to the bifunctional AmiA_Cp, highlighting the different strategies these closely related human pathogens employ in PGN remodeling processes. We used CRISPRi methods to induce conditional genetic knockdown of AmiA_Ct and assessed the effect of knockdown by fluorescence microscopy, revealing that AmiA plays a role in chlamydial cell division earlier than what was expected. Studying cell division processes in intracellular *Chlamydia* spp. may provide novel insights on the broader mechanisms of bacterial cell division.

## Results

### *Chlamydia trachomatis* encodes an AmiA homolog

In a bioinformatic approach, *ct268* from *C. trachomatis* serovar D (*ctl0520* in serovar L2) was found to encode a protein with 59% amino acid sequence identity to cell division amidase AmiA_Cp from *C. pneumoniae* and 36% identity compared to AmiA of *E. coli* (AmiA_Ec). The protein was presumed to be an AmiA homolog and is hereafter referred to as AmiA_Ct. The catalytic site of *E. coli* AmiA is made up of two histidine residues, one aspartic acid residue, and one glutamic acid residue coordinating a zinc atom [35]. An additional glutamic acid is predicted to act as a general base catalyst [36]. *In silico* sequence analysis revealed the active site of AmiA is conserved between *Chlamydia* and *E. coli,* except for D135 and E167 that are not conserved in *Chlamydia* spp. **(Figure 1b)** [31]. Predictions made by Phyre^2^ revealed the active site residues of both chlamydial amidases to be set within a structure resembling a binding grove **(Figure 1a)**. The active center of AmiA_Ec is occluded by a blocking helix that is released upon protein activation [37]. This blocking helix appears to be missing from the chlamydial AmiA orthologs. Activator proteins interact with the interaction helix that induce a conformational switch in the blocking helix, thereby releasing it from the catalytic center. For chlamydial amidases, a regulatory mechanism is still elusive. Canonical helix structures, such as those present on AmiA_Ec are missing on AmiA_Ct and catalytic centers are constantly exposed, hinting at the possibility that chlamydial amidases are active by default **(Figure 1a)**. Besides amidase activity, Klöckner *et al.* demonstrated AmiA_Cp to be bifunctional with additional D,D-carboxypeptidase activity conferred by an SxxK PBP motif around the active serine residue at sequence position 96 [31]. An additional SxN motif is located around S140 (**Figure 1b** and **supplementary figure 1**). AmiA_Ct retains only the SxN PBP motif, which is not associated with PBP activity. *In silico* sequence analysis of amino acid sequences of both chlamydial amidases and AmiA_Ec revealed positional conservation of PBP motifs and active site residues. An exception is E242 in AmiA_Ec, which is shifted towards the distal end of the sequence due to the regulatory domain starting at position S157 (**Figure 1b** and **supplementary figure 1**). As for AmiA_Cp, a periplasmic localization of AmiA_Ct is predicted. While AmiA_Ec is exported to the periplasmic space in a Tat-dependent manner, a Tat system is absent in *Chlamydia*, and AmiA_Cp was found to be secreted Sec-dependently [31]. In line with this, a predicted signal peptide with a cleavage site between positions 47 and 48 is present in AmiA_Ct (**supplementary figure 2**). Thus, our data indicate that *C. trachomatis* harbors an AmiA homolog, which is predicted to be exported to the periplasm.

**Figure 1.**
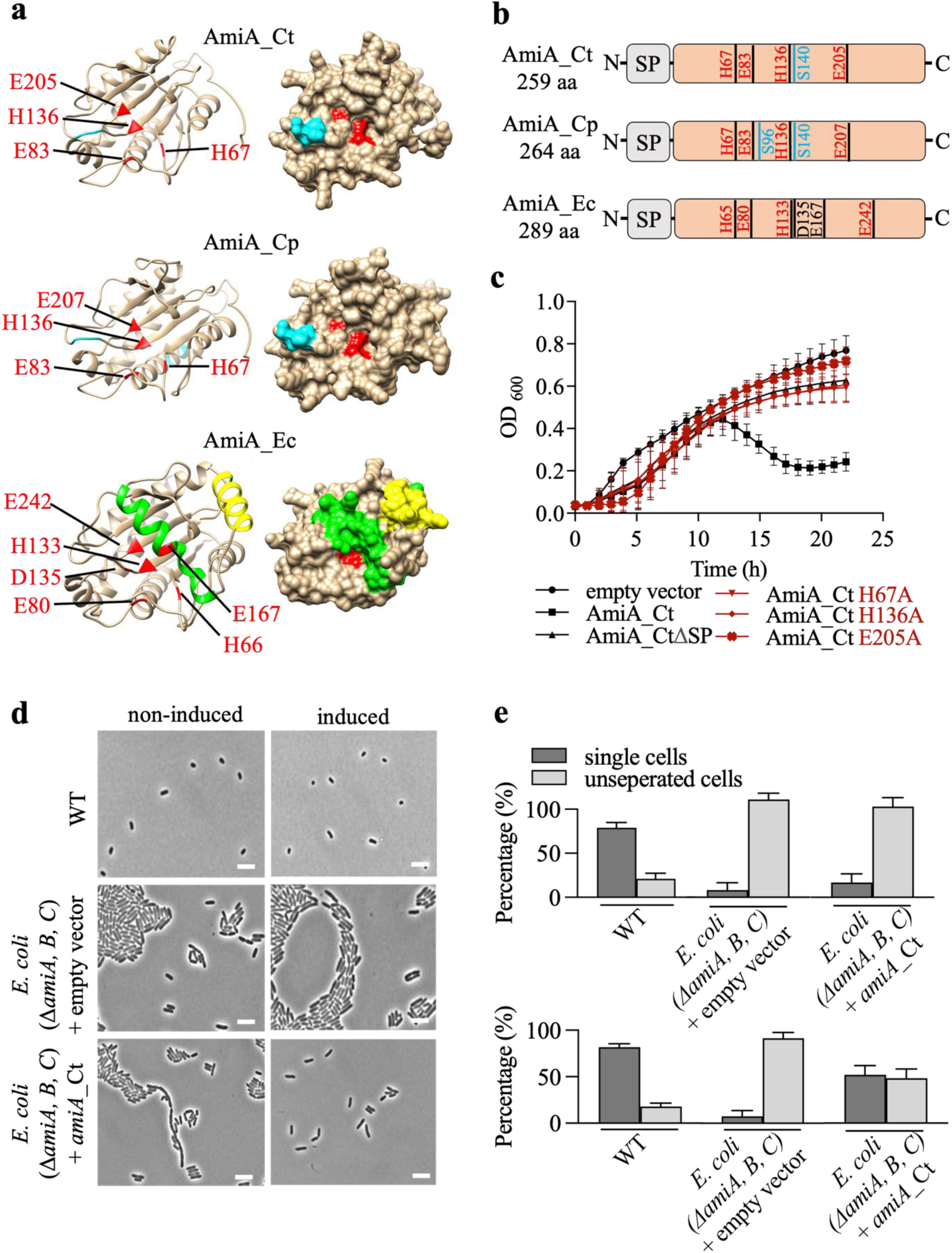
(a) Comparative Phyre2 predictions of AmiA homologs of *C. trachomatis, C. pneumoniae*, and *E. coli* with the residues coordinating the zinc atom within the active site highlighted in red. For AmiA of *E. coli*, the blocking helix is highlighted in green, and the interaction helix is depicted in yellow. The SxN motif in *C. trachomatis* and *C. pneumoniae* and the SxxK motif in *C. pneumoniae* are depicted in blue. A schematic alignment of AmiA homologs **(b)** shows that active site residues (red) are highly conserved, with the exception of D135 and E167 found only in AmiA_Ec. While AmiA_Cp contains two PBP motifs (depicted in blue), AmiA_Ct only retained one PBP motif (SxN) containing the serine residue at position 140. **(c)** Overexpression of *amiA*_Ct lyses the producer strain and shows that both histidine and the glutamic acid residue are equally important for enzymatic activity. Qualitative **(d)** and quantitative **(e)** data reveal that ectopic expression of AmiA_Ct in an amidase-deficient *E. coli* mutant rescues cell separation. Scale bar = 10 µm. Error bars indicate ± s.d. (n = 3).

To determine if AmiA_Ct has amidase activity in *E. coli*, we performed a lysis experiment in *E. coli* in strains overproducing different isoforms of recombinant protein. Upon induction of wild-type AmiA_Ct expression, lysis of the producer strain was observed after 12h with a noticeable drop in OD_600_ **(Figure 1c)**. These results are in line with the results of Klöckner *et al*. for AmiA_Cp [31]. Here, the native signal peptide of AmiA_Ct enabled sufficient periplasmic export without an OmpA leader peptide, and deletion of the predicted signal peptide allowed normal growth of *E. coli* when this isoform was overexpressed in *E. coli*. Mutagenesis of the active site of AmiA_Ct revealed that H67, H136, and E205 (corresponding to H65, H133, and E242 in AmiA_Ec) are equally important for enzymatic activity as mutation of any of these residues rescued growth of *E. coli* in strains overproducing these mutant AmiA_Ct proteins. While the corresponding residue E207 in AmiA_Cp is not required for enzyme function, AmiA_Ec also requires E242 [36]. To see if AmiA_Ct assists cell separation in an amidase deficient *E. coli* mutant (Δ*amiABC*), a complementation assay was performed analogously to the method described by Klöckner *et al*. [31]. Microscopy confirmed that, under uninduced conditions, cells were predominantly present in chains of varying length in the Δ*amiABC* strain. Similar results were obtained for the empty vector control strain under inducing conditions. Induction of AmiA_Ct partially restored the single cell phenotype **(Figure 1d)**. Quantitative analysis revealed >50% single cells under induced conditions compared to <20% single cells in the uninduced population for the Δ*amiABC* strain complemented with AmiA_Ct **(Figure 1e)**. These results confirm that, similar to AmiA from *C. pneumoniae*, AmiA_Ct is functional in free-living bacteria and likely degrades PGN to assist cell separation.

### AmiA_Ct has amidase activity on PGN and precursor molecule lipid II

To test the *in vitro* function of purified AmiA_Ct, we performed dye release assays using Remazol Brilliant Blue-stained PGN as a substrate with recombinant protein. *In vitro* PGN cleavage was previously shown for AmiA_Ec and AmiA_Cp [30, 31], and the latter was used as a control here with activity on par with lysozyme **(positive control; Figure 2a)**. Consistent with its predicted activity, AmiA_Ct hydrolyzed PGN *in vitro* while a purified active site mutant did not **(Figure 2a)**. Reaction efficiency showed pH-dependency, with the highest enzymatic activity in alkaline conditions at pH 8.5 and pH 9.5 **(Figure 2b)**. Additionally, the activity of wild-type or active site mutant H67A AmiA_Ct on PGN precursor lipid II *m*DAP was tested. TLC-based analysis combined with mass spectrometry revealed monofunctional activity on lipid II mDAP that requires an intact active site, releasing the peptide side chain from undecaprenyl-pyrophosphoryl-MurNAc-GlcNAc **(Figure 2c-3)**. AmiA_Ct does not show carboxypeptidase activity like its homolog AmiA_Cp [31]. In conclusion, purified AmiA_Ct hydrolyzes PGN and lipid II *m*DAP *in vitro* and is a monofunctional amidase.

**Figure 2.**
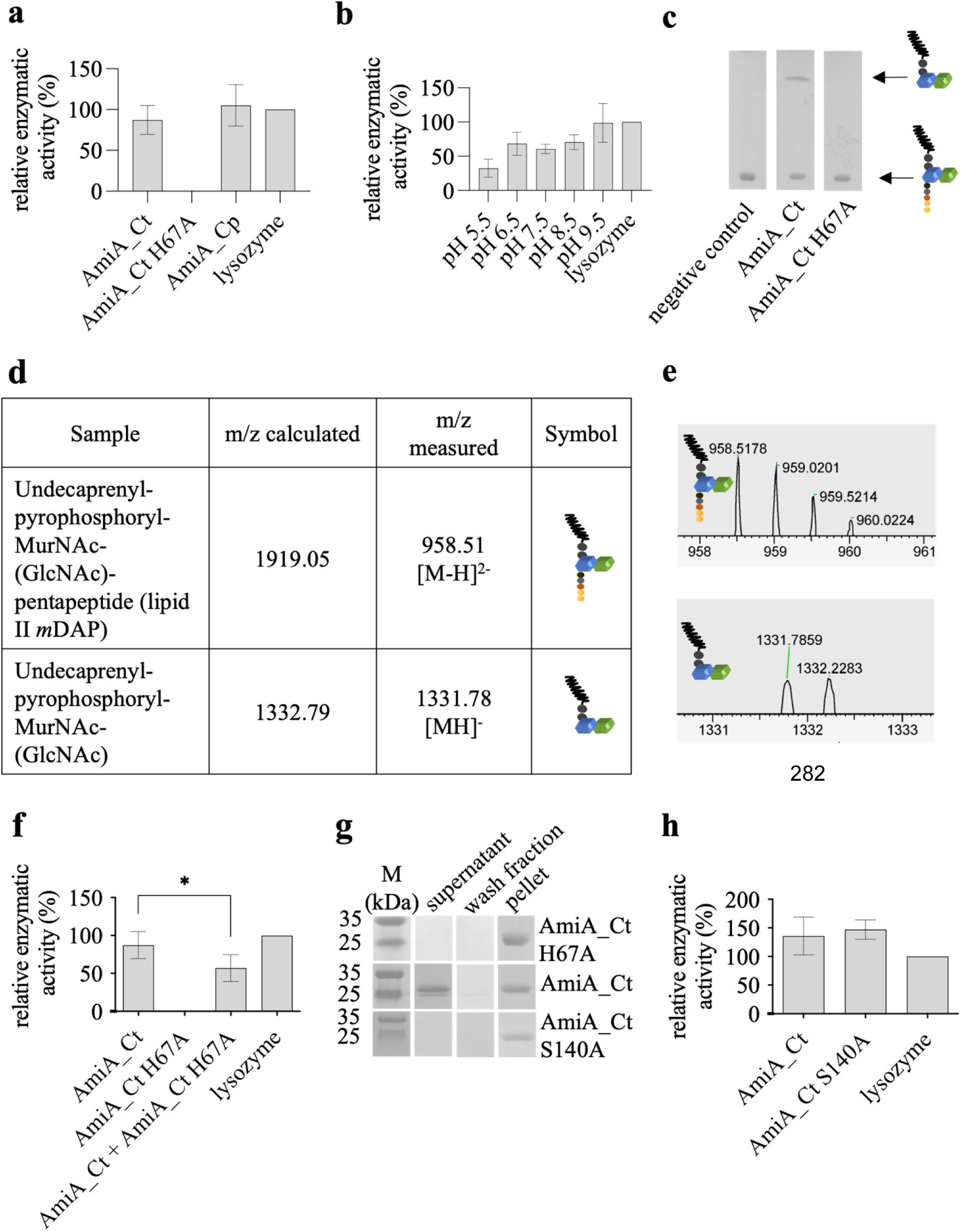
Dye release assays using Remazol Brilliant Blue-stained PGN were performed to assess amidase activity and could show that AmiA_Ct cleaves PGN *in vitro* **(a)**. Enzymatic activity is pH-dependent **(b),** comparable to AmiA_Cp activity on PGN and could be abolished upon mutagenization of the active site **(a)**. TLC **(c)** and MS analysis of AmiA_Ct reaction products **(d,e)** revealed that AmiA_Ct cleaves PGN precursor lipid II and is a monofunctional amidase. When incubated together with the active site mutant in an equimolar ratio, AmiA_Ct showed reduced activity with PGN as a substrate **(f)**. To test binding of the enzyme to the substrate, pulldown experiments were performed **(g)**. Bands on SDS-PAGE represent AmiA_Ct (24.3 kDa). Results showed that PGN binding is independent from enzymatic activity, but is not caused by the rudimentary PBP motif S140. Error bars indicate ± s.d. (n = 3). Ordinary One-way ANOVA * = p ≤ 0.05.

**Figure 3.**
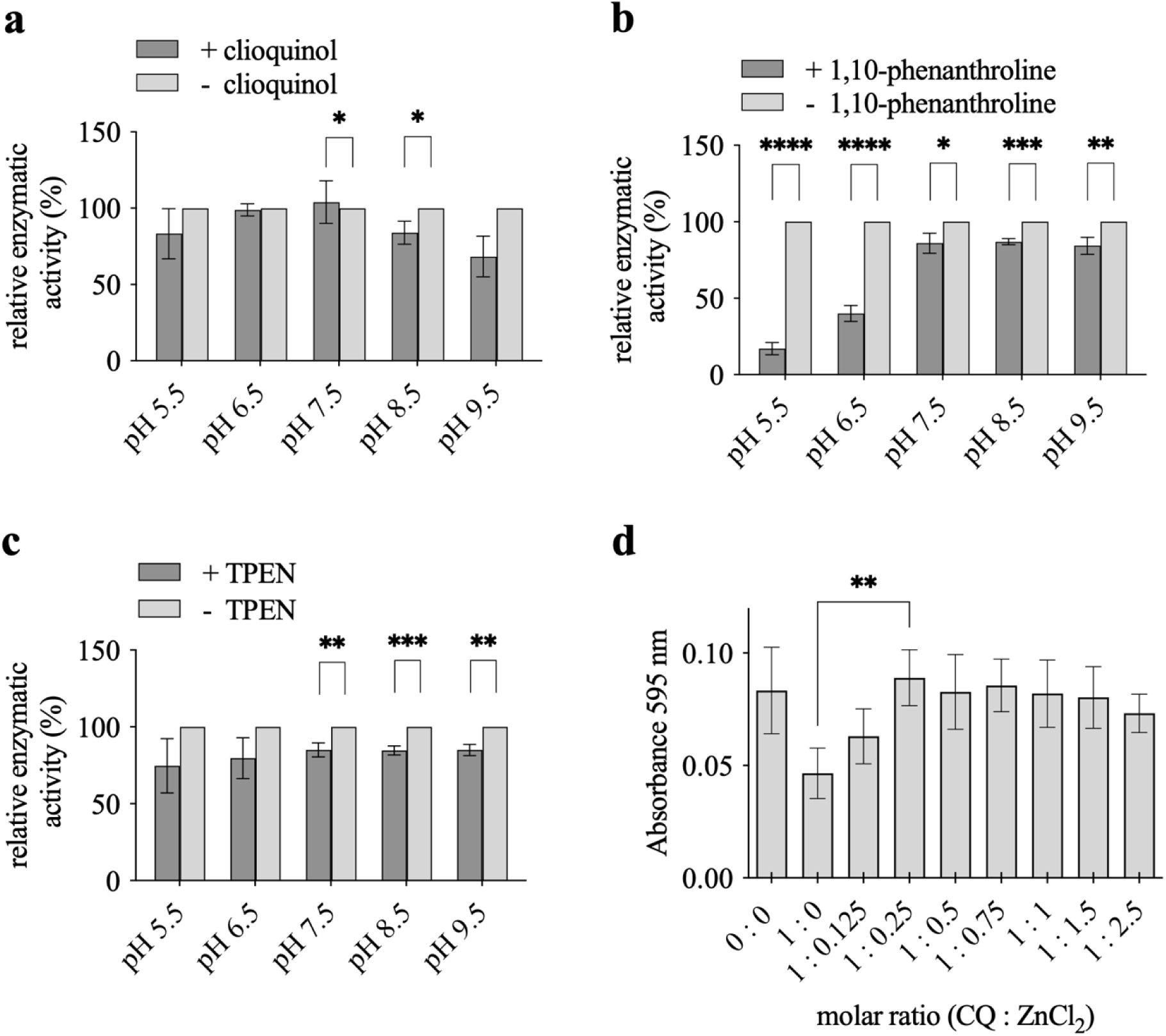
AmiA_Ct is inhibited by chelators of bivalent cations. The antibiotic clioquinol (CQ) **(a)**, 1,10-phenanthroline **(b)** and TPEN **(c)** inhibited AmiA_Ct in dye release assays in a pH-dependent manner. **(d)** Upon addition of excess ZnCl_2_ to the reaction, the inhibitory effect of CQ was prevented, showing that enzyme inhibition relies on the sequestration of the zinc atom by chelators. Error bars indicate ± s.d. (n = 3). * = p ≤ 0.05, ** = p ≤ 0.01, *** = p ≤ 0.001, **** = p ≤ 0.0001.

### PGN binding of AmiA_Ct is independent from enzymatic activity

In PGN dye release assays, as expected, wild-type AmiA_Ct activity approached the lysozyme control whereas the H67A mutant had no detectable activity. AmiA_Ct and the active site mutant AmiA_Ct H67A were also incubated in equimolar amounts and tested. Interestingly, enzymatic activity of the mixed isoforms was significantly lower than the reaction that included only AmiA_Ct **(Figure 2f)**. This raised the question whether a PGN binding motif is present on AmiA_Ct independent of its catalytic site. Bioinformatic analysis of the amino acid sequence did not reveal a putative PGN binding domain. To test binding conditions of AmiA_Ct to PGN, pulldown assays were performed with purified protein and *B. subtilis* PGN as a substrate. Analysis by SDS-PAGE showed that AmiA_Ct H67A bound PGN **(Figure 2g)**. In contrast, native AmiA_Ct bound partly to the substrate, which was also present in the supernatant, indicating that only native AmiA_Ct is able to release the substrate. AmiA_Ct has a PBP motif (SxN) in its sequence, which does not comprise any PBP-related activity. To test whether it mediates PGN binding, we mutagenized AmiA_Ct by replacing residue S140 with alanine (S140A). Pulldown assays were performed with the purified protein, which showed its complete binding to PGN, similar to the H67A mutant. However, in dye release assays, the protein was fully functional as an amidase, indicating that the catalytic center was not affected **(Figure 2h)**. Our data indicate that PGN binding of AmiA_Ct is independent from its amidase activity.

### Amidase activity/AmiA_Ct can be inhibited by chelators of bivalent cations

As *in vitro* activity of AmiA_Ct was demonstrated in dye release assays, we next tested whether we could inhibit the *in vitro* reaction. A common characteristic for the amidase_3 family as classified by Pfam is the coordination of a zinc atom in the catalytic center. Hence, cell division amidases are zinc-dependent enzymes. Hypothetically, chelators of bivalent cations should sequester the zinc atom from the active center and thereby inactivate AmiA_Ct in a dye release assay. We tested different chelator antibiotics under similar conditions as described in **Figure 2**. The antibiotic clioquinol modestly but significantly inhibited the reaction at pH 7.5 and 8.5 with a further, not statistically significant reduction, at pH 9.5 **(Figure 3a)**. Inhibition by 1,10-phenanthroline was detectable at all pH values tested, with a dramatic reduction in activity at acidic conditions around pH 5.5 and pH 6.5 **(Figure 3b)**. TPEN showed modest but significant inhibition at neutral and alkaline pH **(Figure 3c)**. To clarify if inhibition is reversible, a dye release assay including clioquinol and zinc chloride in ascending molar ratios was performed. Enzymatic activity of AmiA_Ct was restored at a molar ratio of 1 : 0.25 (clioquinol : ZnCl_2_) **(Figure 3d)**. Taken together, these data indicate the *in vitro* hydrolytic activity of AmiA_Ct can be inhibited by metal chelators in a pH-dependent manner.

### CRISPRi-mediated knockdown of *amiA* alters chlamydial morphology and negatively affects generation of infectious progeny

The role AmiA plays in chlamydial cell division processes was further investigated by targeting *amiA*_Ct utilizing a CRISPRi-mediated knockdown (KD) approach. A CRISPRi vector targeting an intragenic region of *amiA*_Ct (pLCRia (*amiA*); *amiA* KD) was transformed into *C. trachomatis*. This system relies on the constitutive expression of a guide RNA (gRNA) and anhydrotetracycline (aTc)-inducible expression of a catalytically dead dCas9 [38]. As a control, we compared effects of knockdown to a strain carrying a vector encoding a non-targeting gRNA with dCas9 (pLCRia (NT); NT). We also generated a complementing vector wherein an *amiA_6xH* allele is transcriptionally fused to the dCas9 gene; a strain carrying this complementing vector was also generated (pLCRia-*amiA_6xH* (*amiA*); *amiA* KDcomp). Therefore, this will restore *amiA* expression when the chromosomal copy is transcriptionally repressed [39].

HeLa cells were infected with each of the three strains, and dCas9 expression was induced or not at 10hpi with 5nM aTc. Total RNA and genomic DNA (gDNA) were collected at 10, 14, and 24hpi from all samples. Overall, all strains and conditions showed similar levels of genomic DNA at each timepoint, indicating that dCas9 expression did not disrupt DNA replication **(Figure 4a)**. Transcript levels for *amiA*, *murE* (encoded 5’ to *amiA* in an operon), *euo* (an early cycle gene), and *omcB* (a late cycle gene) were measured by RT-qPCR and normalized to gDNA. For each of the conditions and strains tested **(Figure 4b-d)**, there were no significant changes in transcript levels for *murE*, *euo*, or *omcB*. However, *amiA* transcripts were significantly reduced in the *amiA* KD strain after inducing dCas9 expression, indicating successful knockdown of this target. In the *amiA* KDcomp strain, *amiA* transcripts were elevated in uninduced conditions approximately 10-fold above the uninduced conditions in the NT and *amiA* KD strains. This suggests an internal promoter in the dCas9 gene may drive expression of *amiA_6xH*. Nonetheless, after inducing dCas9 expression, *amiA* transcripts returned to a wild-type level (∼0.01ng), consistent with the intragenic target for the *amiA* gRNA. Overall, these data indicate (i) successful knockdown of *amiA* and (ii) complementation in the *amiA* KDcomp strain.

**Figure 4:**
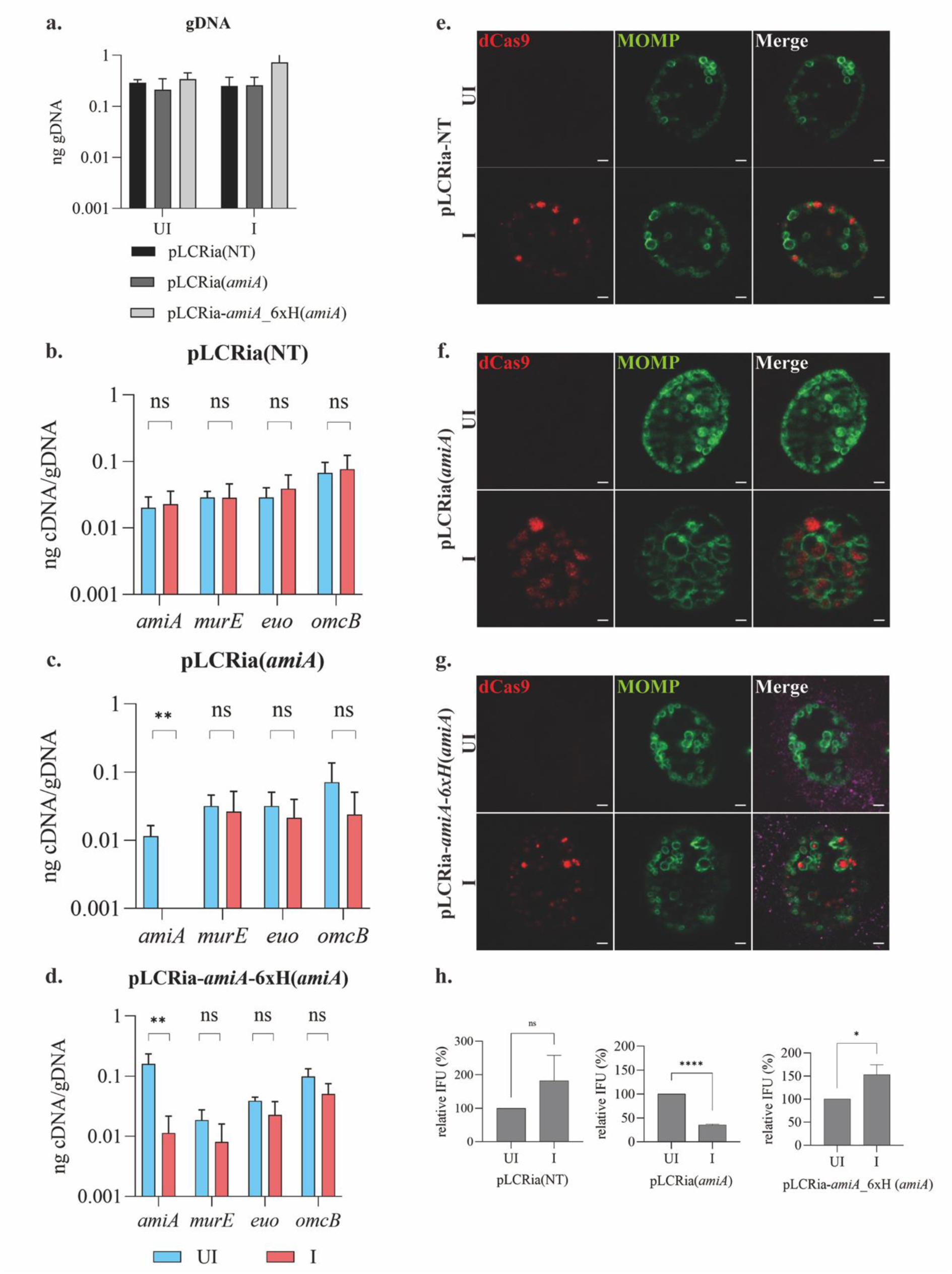
CRISPRi-mediated knockdown of *amiA*_Ct negatively impacts chlamydial morphology and infectious progeny production. Constructs encoding dCas9-gRNA CRISPRi systems targeting *amiA*_Ct (pLCRia(*amiA*)), targeting *amiA*_Ct while overexpressing *amiA*_Ct (pLCRia-amiA_6xH(*amiA*)), or a non-targeting gRNA (pLCRia(NT)) were transformed into *C. trachomatis* (-pL2). HeLa cells were infected with the transformants, and expression of the CRISPRi system was induced or not with 5 nM aTc at 10 hpi. At 24 hpi, RNA and genomic DNA were isolated and used for (RT)-qPCR to measure transcript and genomic DNA levels, coverslips were fixed for immunofluorescence analysis (IFA), or samples were collected for inclusion forming unit (IFU) assays. **(a)** Levels of genomic DNA of pLCRia(NT), pLCRia(*amiA*), and pLCRia-amiA_6xH(*amiA*) strains at 24 hpi in uninduced (UI) and induced (I) conditions. **(b-d)** Transcript levels of *amiA*, *murE*, *euo*, and *omcB* in dCas9-uninduced and induced conditions at 24 hpi in the pLCRia(NT) **(b)**, pLCRia(*amiA*) **(c)**, and pLCRia-amiA_6xH(*amiA*) strains **(d)**. (**e-g**) IFA images in dCas9-uninduced and induced conditions at 24 hpi in the pLCRia(NT) (**e**), pLCRia(*amiA*) (**f**), and pLCRia-amiA_6xH(*amiA*) (**g**) strains. For IFA controls, the major outer membrane protein (MOMP in green) and dCas9 (red) were labelled. Images were acquired on a Zeiss AxioImager.Z2 equipped with an Apotome2 using a 100x objective. Error bars represent ± s.d. (n=3). Unpaired t-test, two-tailed. * p ≤ 0.05, ** p ≤ 0.01. Scale bar = 2 µm if not indicated otherwise. (**h**). IFU assay to measure effect of knockdown on infectious progeny production. Samples were collected for each condition in each strain and titrated onto a fresh monolayer of cells to quantify the number of inclusions derived from infectious EBs produced during the primary infection. * p < 0.05, **** p < 0.0001, ns = not significant by unpaired t-test, two-tailed.

We next assessed the broad morphologic effects of *amiA* knockdown by immunofluorescence analysis (IFA) and quantified the ability of *Chlamydia* to complete its developmental cycle by measuring inclusion forming units (IFUs) – a proxy for infectious EBs. For each of these samples, HeLa cells were infected with the three strains, and dCas9 expression was induced or not as above. At 24hpi IFA samples were fixed and processed, and IFU samples were collected. IFA samples were labeled with antibodies recognizing the chlamydial major outer membrane protein (MOMP) and dCas9. Typical chlamydial morphologies (a mixture of rings of different sizes representing individual bacteria) were observed by IFA for the NT strain under both inducing and uninducing conditions with dCas9 labeling evident in the inducing conditions **(Figure 4e)**. For the *<u>amiA</u>* KD strain, organism morphology appeared normal in the uninducing conditions. However, after inducing the *amiA* KD strain, enlarged RBs were evident **(Figure 4d)**, consistent with a block in cell division as has been observed with beta-lactam treatment [40]. The abnormal morphology was not evident in the *amiA* KDcomp strain under the conditions tested **(Figure 4f)**. Chlamydial chromosomal copy number was not significantly affected under induction for any of the three strains observed **(Figure 4g)**, however, IFU measurements indicated a statistically significant reduction in IFUs during *amiA* knockdown that was not detected in the NT or *amiA* KDcomp strains **(Figure 4h)**. Collectively, these data indicate that *amiA* knockdown has a negative impact on chlamydial morphology that results in a reduction in IFU production.

### CRISPRi-mediated knockdown of *amiA* induces an inhibited cell division phenotype in *Chlamydia* and alters PGN labeling

A ‘clickable’ D-alanine dipeptide (EDA-DA) was used to detect chlamydial PGN while immunostaining against chlamydial MOMP to enable the visualization of chlamydial cells. When the effects of *amiA* knockdown were monitored at 24 hpi, chlamydial cells appeared enlarged and displayed a phenotype indicative of inhibited cell division, as shown by MOMP-directed immunostaining **(Figure 5a)**. Parent and daughter cells were clearly visible, however, they exhibited significant size asymmetry, consistent with a pause during the budding phase of the pathogen’s unique division process [13, 24, 25, 41]. In a complementation approach, a CRISPRi vector inducing *amiA* knockdown while simultaneously overexpressing *amiA*_Ct was transformed into *C. trachomatis*. Upon induction, complementation resulted in normal chlamydial development **(Figure 5b)**. PGN was visualized via click chemistry and revealed effects of *amiA* knockdown on chlamydial PGN, and it was found to localize in discrete patched areas proximal to budding daughter cells. Upon complementation, PGN rings with septal localization were reestablished. When these three *<u>C. trachomatis</u>* strains were allowed to grow normally for 23 hours and then induced for 1 hour, the *amiA* knockdown strain exhibited higher intensity PGN labeling than either the non-targeting control strain (NT) or the complemented strain **(Figure 5c)**. Quantitative analysis showed significantly enlarged PGN object volumes in the induced knockdown strain compared to the uninduced strain. Similar results were obtained for integrated density, which is a product of PGN volume and the mean intensity of labeled PGN **(Figure 5d)**.

**Figure 5.**
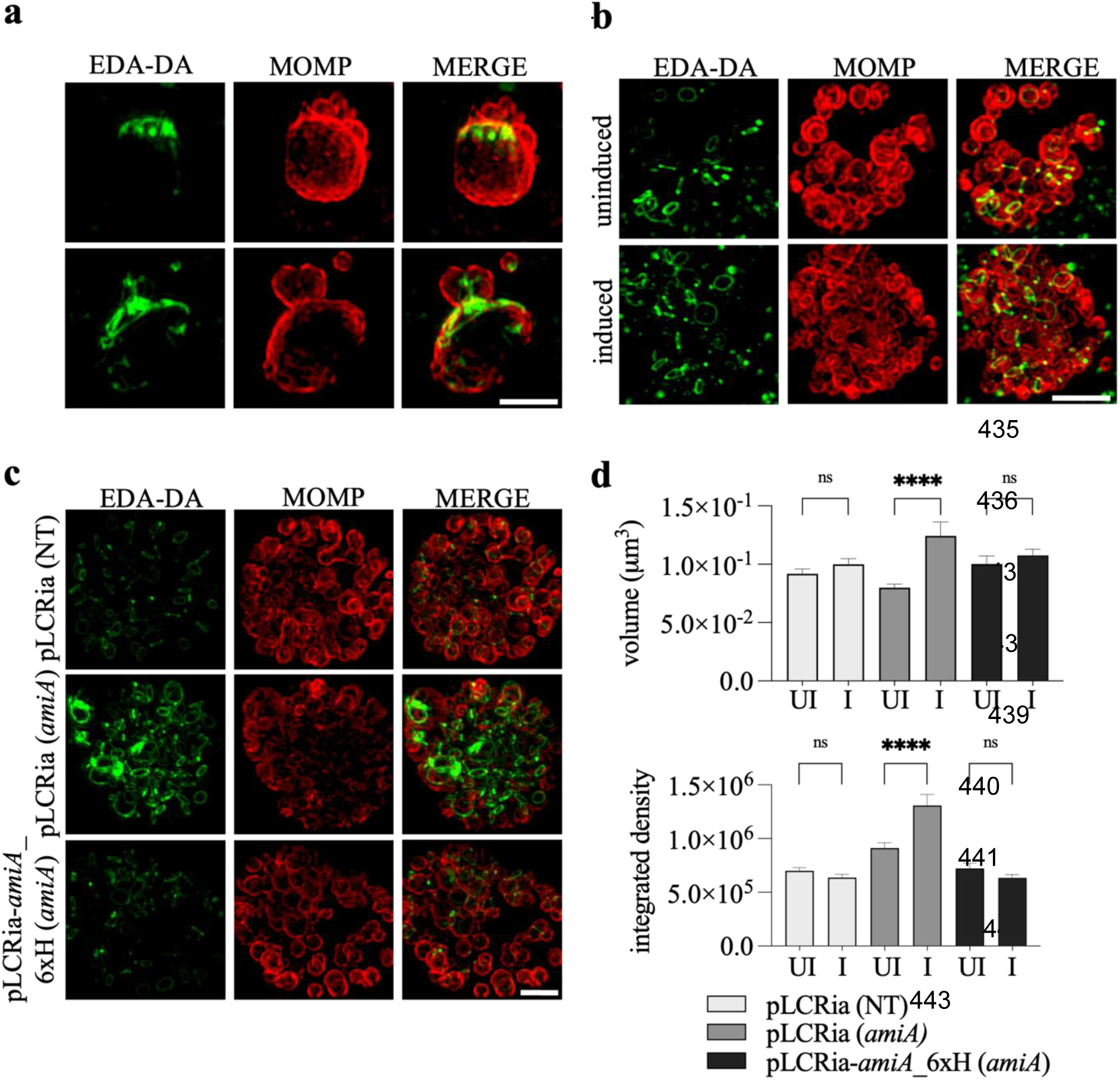
Determination of the effects of CRISPRi-mediated knockdown of AmiA_Ct. Constructs encoding a dCas9-gRNA CRISPRi system targeting *amiA*_Ct (pLCRia (*amiA*)) **(a)** and one simultaneously overexpressing *amiA*_Ct (pLCRia-*amiA*_6xH (*amiA*)) **(b)** were transformed into *C. trachomatis* (-pL2). McCoy cells were infected with these transformants and expression of the CRISPRi system was induced with 1 nM aTc at 2 hpi. Clickable D-alanine dipeptides (EDA-DA) were used for PGN labeling. Cells were fixed 24 hpi and imaged. Knockdown of *amiA*_Ct resulted in enlarged chlamydial cells with impaired cell division halted in the budding phase, and PGN concentrated to the budding poles **(a)**. Complementation of *amiA*_Ct restored normal chlamydial phenotypes and PGN localization **(b)**. When induced at 23 hpi with 5 nM aTc, enhanced PGN labeling of the *amiA*-targeting CRISPRi-system is revealed compared to the complemented and the non-targeting (NT) construct **(c)**, which was shown quantitatively in **(d)**. Values in **(d)** are presented in arbitrary fluorescence units. MOMP: major outer membrane protein. NT: non-targeting. Unpaired t-test, two-tailed: * p ≤ 0.05, ** p ≤ 0.01, *** p ≤ 0.001, **** p ≤ 0.0001. Error bars indicate ± s.d. (n=3). Scale bar = 2 µm.

## Discussion

Pathogenic members of the Chlamydiae synthesize a transient PGN ring during their unique FtsZ-independent polarized budding division process [12, 13, 25]. The PGN ring requires constant remodeling to dilate and constrict, and PGN is degraded after cell division and recycled for future division events. In this study we show that AmiA_Ct, the only annotated cell division amidase retained in the genetically streamlined *Chlamydia trachomatis*, is a monofunctional amidase that can bind and degrade PGN, can complement an *E. coli* Δ*amiABC* mutant, and is a critical regulator of chlamydial cell division.

A major distinction between cell division amidases in free-living *E. coli* and intracellular *Chlamydia* is the apparent lack of a regulatory mechanism in chlamydial amidase orthologs. According to *in silico* structural models of chlamydial amidases, the enzyme active sites are constantly exposed, suggesting that, as soon as AmiA is exported to the periplasm and contacts PGN, it will degrade it. However, this model would lead to additional questions. For example, is AmiA degraded after cell division is completed and resynthesized during the RB cell cycle? If not, and AmiA is present in the periplasm during the developmental cycle, then how is it prevented from degrading the growing PGN ring during chlamydial division? In *E. coli* cell division amidases adopt a resting state with an autoinhibited conformation, in which the active sites are occluded by a blocking helix [35, 37]. Interaction with LytM domain-containing proteins EnvC, in the case of AmiA and AmiB, and NlpD, for AmiC, leads to withdrawal of the blocking helix from the active site [30]. While the majority of LytM proteins are metalloendopeptidases, *E. coli* homologs are among the few exceptions that do not possess hydrolytic activity [30]. *E. coli* deletion mutants of NlpD and EnvC lead to the formation of cell chains, similar to amidase deletion mutants [42]. Like cell division amidases in *E. coli*, EnvC and likely NlpD are not active by default but display a similar autoinhibition mechanism. A domain referred to as the restraining arm occludes the LytM domain binding grove, intended for an interaction helix, part of the amidase regulatory domain, to bind [35, 43]. According to a model by Yang *et al.*, the restraining arm is removed from the binding groove of LytM proteins by interaction with FtsEX, which has a conserved role in bacterial cell separation, as shown for example for *S. pneumoniae* [44]. Mutants display comparable cell division defects and phenotypes to what is observed in amidase and EnvC/NlpD deletion mutants [45]. ATP hydrolysis by ATPase FtsE in the cytoplasm is proposed to cause transmembrane protein FtsX to trigger a conformational change in the periplasmic LytM-domain proteins, retrieving the restraining arm from the LytM binding grove and thereby facilitating amidase activation by these enzymes [45]. In chlamydiae, homologs of LytM proteins EnvC or NlpD have not been retained nor do they encode FtsEX homologs to initiate the activation cascade. Previously, we showed that *C. pneumoniae* protein Cpn0902 has been misannotated as NlpD since it confers carboxypeptidase activity and thereby likely participates in chlamydial PGN remodeling [31].

Cook *et al.* propose the active site of AmiA in *E. coli* to be composed of two histidine residues (H65, H133), one glutamic acid residue (E80) and D135 and E167. While E167 is predicted to be part of the blocking helix, all five residues are described to be involved in coordination of zinc in the enzyme’s active site [35]. The authors propose D167 to serve as a latch holding the blocking helix in place in the autoinhibited conformation. As previously proposed by our group, all corresponding residues except D135 and E167 have been positionally conserved in chlamydial amidases [31]. We propose that like in the *E. coli* homolog, these residues are involved in zinc coordination. However, the requirement of E205 of AmiA_Ct for coordination can only be hypothesized, as the corresponding residue in *E. coli* AmiA, E242, is suggested to act as a base catalyst [36]. As no crystal structures of chlamydial amidases are available, the coordination status of zinc within the active site cannot be completely clarified to date. However, *in vitro*, mutation of H67 in purified AmiA_Ct led to a full loss of enzymatic activity in dye release assays, reinforcing the residue’s function as part of the active center coordinating the catalytic zinc. In a lysis experiment, amino acid substitutions of both histidine residues and E205 in heterologously overproduced AmiA_Ct led to a complete loss of function of the protein. This indicates that each of these three residues is equally important for enzymatic activity of AmiA_Ct. Conversely, the corresponding residue to E205 in AmiA_Cp, E207, was not needed for lysis of the *E. coli* producer strain in a similar experiment we performed previously, indicating a differential architecture of the catalytic centers of these closely related proteins [31]. As the substitution of E80 in *E. coli* AmiA did not lead to a full loss of *in vitro* activity in previous experiments [36], a chlamydial AmiA mutant with a positionally conserved substitution was not included in this experiment.

Like AmiA_Cp, the homolog from *C. trachomatis* degraded PGN *in vitro* [31]. In general, the results from our experiments using both PGN and lipid II as substrates further demonstrate that chlamydial amidases are active by default and do not require an additional protein partner for activation. However, the binding mode of these enzymes to PGN is unknown. Interestingly, of the three periplasmic cell division amidases in *E. coli*, AmiA is the smallest. The enzyme consists of a single globular domain and lacks, unlike AmiB and AmiC in *E. coli*, an N-terminal AmiN-domain conferring binding affinity towards PGN [46]. AmiA homologs in *C. pneumoniae* and *C. trachomatis* also do not contain AmiN-domains, and bioinformatic searches with the amino acid sequences also did not reveal the presence of any known PGN binding domains previously identified in hydrolases such as SPOR, CHAPS, SHb3, or LysM. Zoll *et al*. investigated substrate binding of AmiE, a subunit of autolysin AtlA in *Staphylococcus epidermidis* that also coordinates a zinc atom within its catalytic core. Upon mutagenesis of active site residues, the enzyme was catalytically inactive. However, PGN binding was not impaired by catalytic inactivation but instead had a positive effect on substrate affinity. One of the active site residues, H177, is not thought to be involved in zinc coordination but rather in the stabilization of the enzyme-substrate transition state during the catalytic process. Proposed to enable a water-mediated nucleophilic attack, a stabilizing function of the transition state within the catalytic process is attributed to the enzyme [47, 48]. The authors suggest this increase in binding affinity to be due to a smaller structure resulting from the replacement of histidine by alanine. In pulldown experiments, binding affinity of purified AmiA_Ct and active site mutant AmiA_Ct H67A towards PGN was assessed and showed that the catalytically inactive mutant bound to PGN as a substrate completely, whereas the active enzyme had reduced binding affinity. In analogy to the hypothesis suggested by Zoll *et al*., replacement of histidine by alanine might result in a smaller structure that stabilizes the transition state of catalysis and might lead to permanent binding between enzyme and substrate. Replacements of highly conserved residues of an enzyme’s catalytic center could also lead to the construction of a substrate trap, as described by Flint *et al.* for protein tyrosine phosphatases, serving as a possible explanation for higher PGN binding affinity of a catalytically inactive enzyme [48]. Also, replacement of histidine by alanine changes physicochemical properties of the enzyme’s catalytic core, and these effects influencing binding affinities towards a substrate could be possible. Processes of substrate binding, catalysis and release by an enzyme are highly dynamic, possibly explaining why AmiA_Ct is only bound partially to the substrate, as the experiment only represents limited time points. Our results indicate that PGN binding is independent from enzymatic activity, but a specific region of the protein conferring binding affinity to the substrate could not be identified. Testing if replacement of active site residues is mediating a direct increase in binding affinity, and which regions of the protein are involved, is yet to be investigated.

Although the mode of substrate recognition and binding remains to be fully clarified in chlamydial amidases, the coordinated zinc in the active center of the enzyme is conserved across this enzyme class and most likely plays an important role. To investigate further, experiments with compounds known to chelate bivalent cations with a that have been administrated in *in vitro* and *in vivo* settings were performed. 1,10-phenanthroline is a membrane-permeable compound and a chelator of bivalent cations. Antibiotic action against some Gram-positive, Gram-negative and acid-fast bacteria has been demonstrated, and antifungal activity has also been documented [49]. *In vitro*, the compound caused inhibition of pancreatic carboxypeptidase A by sequestration of zinc from the enzyme’s active center [50, 51]. AmiC of *Neisseria gonorrhoeae*, the only cell division amidase retained in the organism, showed a decrease of *in vitro* activity upon 1,10-phenanthroline treatment at pH 8.5 [52]. Moreover, a purified amidase of *Ochrobactrum anthropi* was inhibited by 1,10-phenanthroline at pH 8.0 and activity could be restored by adding zinc, manganese or magnesium to the reaction [53]. Supply of other transition metals than zinc is thought to mediate release of previously chelator-bound zinc, which can then be bound by an apoenzyme. While studying the substrate spectrum of AmiA of *E. coli*, Lupoli *et al*. successfully inhibited the *in vitro* reaction by application of 1,10-phenanthroline at pH 7.5 [36, 54]. We demonstrated pH-dependent inhibition of AmiA_Ct by 1,10-phenanthroline, Tetrakis(2-pyridylmethyl)ethylenediamine (TPEN), and by the antibiotic clioquinol. Generally, metal complex formation depends on pH conditions, as known for EDTA, which has increased affinity for metal complexes with increasing pH values [54]. The antibiotic clioquinol was also included as a chelator of bivalent cations. Formerly, clioquinol was used to treat severe intestinal diseases such as lambliasis, shigellosis, or amoebal infections of the intestinal tract, and it is still in use as a topic antiseptic in mild skin infections [55]. As shown for 1,10-phenanthroline, inhibition by clioquinol is pH dependent and most effective in neutral to alkaline conditions of pH 7.5 – 8.5. As Sonke *et al.* showed for an amidase of *Ochrobactrum anthropi*, in this experiment, AmiA_Ct activity could be reinstated by supply of additional zinc to the reaction, highlighting the fact that inhibition is caused by zinc sequestration [53]. TPEN is a specific zinc chelator and is also suggested to have antifungal and antibacterial activity. In carbapenem-resistant *Enterobacteriaceae*, reduction of *in vitro* activity of metal-dependent carbapenemase has been observed, possibly conferring increased sensitivity towards meropenem treatment [56]. TPEN significantly inhibited AmiA_Ct *in vitro* at an alkaline pH. The tested compounds represent an opportunity to further characterize the function of chlamydial amidases, and experiments with *E. coli* AmiA could also be considered in the future.

Besides a putative SpoIID homolog acting as a lytic glycosyltransferase [29], as shown for the *Chlamydia*-related bacterium *Waddlia chondrophila*, AmiA is the only known PGN-degrading enzyme in *Chlamydia* spp. to date. Using CRISPRi knockdown technology [38, 57], effects of genetic knockdown of *amiA*_Ct were investigated. Our results revealed that, under *amiA* knockdown conditions, cells displayed a phenotype similar to effects of inhibiting cell division (e.g., with penicillin or D-cycloserine [58]). Knockdown also affected PGN, as it accumulated at the budding poles of cells with inhibited cell division phenotypes. Both regular cell morphology and PGN localization could be restored upon complementation of knockdown with an additional allele of *amiA*, confirming that the observed effects are caused by *amiA* knockdown. Upon knockdown of *amiA*, the degradation of PGN by *Chlamydia* is likely to be significantly impaired while PGN biosynthesis is still functional. This would lead to an accumulation of PGN. Liechti *et al*. hypothesized that altered morphology in *Chlamydia* is a result of impaired cell division due to inhibition of PGN synthesis or failure of the PGN ring to assemble [59]. Previously, we suggested a central function of AmiA within an interconnected cycle of PGN biosynthesis, processing, and recycling of PGN [33]. Therefore, the absence or inactivity of chlamydial AmiA, the only cell division amidase of the organism, could lead to disruption in PGN metabolism. This is consistent with increased PGN volumes that have been measured during *amiA* knockdown. According to Liechti (2021), PGN object volumes are a result of the balance between catabolic and anabolic PGN metabolism [24]. Upon shift of the balance to either side between these processes, PGN object volumes and sizes are likely to decrease or increase. Therefore, increased PGN labeling intensity is suggested to be a result of a high level of PGN synthesis outweighing catabolic processes. As *Chlamydia* is unable to degrade the PGN ring during its cell cycle under *amiA* knockdown conditions, this will lead to a block in cell division and a concomitant reduction in production of infectious EBs as we demonstrated. Interestingly, our observations also indicate that AmiA-dependent PG turnover in *C. trachomatis* appears to be essential not only for septation, but also the expansion of the PGN ring that occurs during the budding phase of the chlamydial division process [24, 41].

Overall, our data highlight a critical function for the AmiA amidase in chlamydial cell division and chlamydial growth more generally. Further work is necessary to understand how AmiA function is regulated in *Chlamydia* as well as the interplay between PGN synthesis and degradation during division.

## Materials and Methods

### Bacterial strains, tissue culture, plasmids, primers and growth conditions

Bacterial strains and plasmids used in this work are listed in tables 1 and 2. Cells were grown in lysogeny broth (LB) (10 g/L tryptone, 5 g/L yeast extract and 10 g/L NaCl) at 30 °C or 37 °C at 120 rpm. Antibiotics were added to the autoclaved medium in final concentrations of 34 µg/mL or 100 µg/mL for chloramphenicol and ampicillin, respectively. The human epithelial cell line HeLa was used for inclusion-forming-unit (IFU) assays, gDNA and RNA extraction. Mouse fibroblast cell line McCoy was used for chlamydial transformation. Cell lines were routinely cultured in Dulbecco’s Modified Eagle Medium (DMEM) + 10 % fetal bovine serum (FBS) + 10 µg/mL gentamycin at 37 °C and 5 % CO_2_. The plasmid-free strain *C. trachomatis* L2 was used for chlamydial transformation. HeLa cells were infected with chlamydial transformants in DMEM + 10 % FBS containing 10 µg/mL gentamycin. EBs of chlamydial transformants were thawed and diluted in HBSS to the indicated MOI and added to the cell monolayer. For synchronous infection, cells were centrifuged at 400 *x g* for 15 min, the incubated at 37 °C for 15 min. After aspiration of inoculum and media exchange, cells were incubated until indicated time.

**Table 1:**
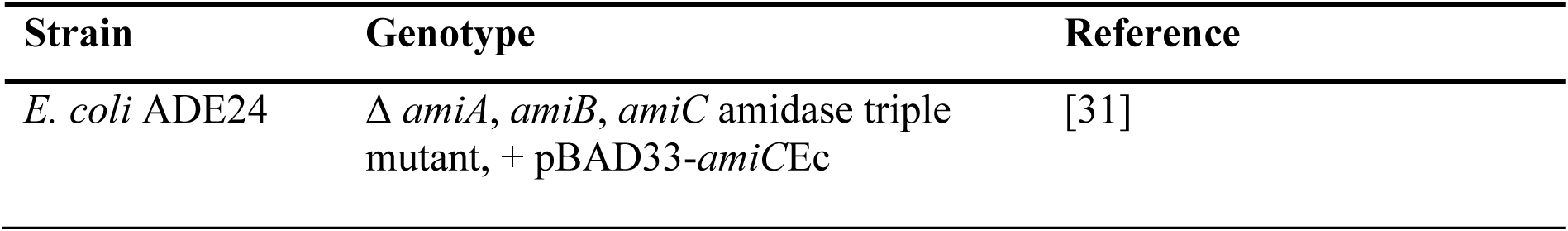
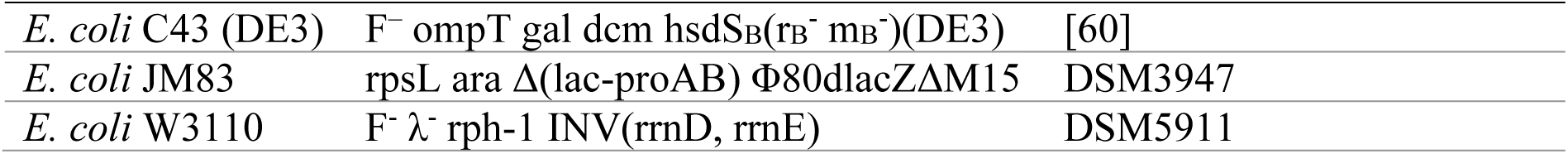
Bacterial strains.

| Strain | Genotype | Reference |
| --- | --- | --- |
| *E. coli* ADE24 | $\Delta$ *amiA*, *amiB*, *amiC* amidase triple mutant, + pBAD33-*amiCEc* | [31] |
| *E. coli* C43 (DE3) | F<sup>-</sup> ompT gal dcm hsdS<sub>B</sub>(rB<sup>-</sup> mB<sup>-</sup>)(DE3) | [60] |
| *E. coli* JM83 | rpsL ara Δ(lac-proAB) Φ80dlacZΔM15 | DSM3947 |
| *E. coli* W3110 | F<sup>-</sup> λ<sup>-</sup> rph-1 INV(rrnD, rrnE) | DSM5911 |

**Table 2:** Plasmids.

| Plasmid | Characteristic | Reference |
| --- | --- | --- |
| pASK-IBA-3 | Expression plasmid, AHT-inducible, C-terminal Strep-tag II, Amp<sup>R</sup> | IBA Lifesciences (Göttingen, Germany) |
| pASK-IBA-3_ct268 | Expression plasmid, AHT-inducible, C-terminal Strep-tag II, Amp<sup>R</sup>, encodes AmiA_Ct | This work |
| pASK-IBA-3C | Expression plasmid, AHT-inducible, C-terminal Strep-tag II, Cam<sup>R</sup> | IBA Lifesciences |
| pASK-3C_ct268 | Expression plasmid, AHT-inducible, C-terminal Strep-tag II, Cam<sup>R</sup>, encodes AmiA_Ct | This work |
| pASK-3C_ct268ΔSP | Expression plasmid, AHT-inducible, C-terminal Strep-tag II, Cam<sup>R</sup>, encodes AmiA_CtΔSP | This work |
| pASK-3C_ct268_H67A | Expression plasmid, AHT-inducible, C-terminal Strep-tag II, Cam<sup>R</sup>, encodes AmiA_Ct_H67A | This work |
| pASK-3C_ct268_H136A | Expression plasmid, AHT-inducible, C-terminal Strep-tag II, Cam<sup>R</sup>, encodes AmiA_Ct_H136A | This work |
| pASK-3C_ct268_E205A | Expression plasmid, AHT-inducible, C-terminal Strep-tag II, Cam<sup>R</sup>, encodes AmiA_Ct_E205A | This work |
| pBOMBL-Sa_dCas9vaa:L2 | dCas9-based conditional knockdown plasmid, AHT-inducible, Amp<sup>R</sup> | [57] |
| pBOMBLCRia(*amiA*) | aTc-inducible CRISPRi-mediated knockdown of *amiA* in *C. trachomatis* L2 | This work |
| pBOMBLCRia-*amiA*_6xH(*amiA*) | aTc-inducible CRISPRi-mediated knockdown of *amiA* in *C. trachomatis* L2, complementation with 6xH-tagged AmiA | This work |
| pBOMBLCRia(NT) | aTc-inducible CRISPRi-mediated knockdown targeted against a | Hatch and Ouellette, 2023 |
|  | random sequence used as a negative control |  |
| pET-52b | Expression plasmid, T7 promotor, autoinducible in (DE3)-strains, N-terminal Strep-tag II, Amp<sup>R</sup> | Novagen ® (Merck, Darmstadt, Germany) |
| pET-52b_*ct268* | Expression plasmid, T7 promotor, autoinducible in (DE3)-strains, N-terminal Strep-tag II, Amp<sup>R</sup>, encodes AmiA Ct | This work |
| pET-52b_*ct268*ΔSP | Expression plasmid, T7 promotor, autoinducible in (DE3)-strains, N-terminal Strep-tag II, Amp<sup>R</sup>, encodes AmiA CtΔSP | This work |
| pET-52b_*ct268*_H67A | Expression plasmid, T7 promotor, autoinducible in (DE3)-strains, N-terminal Strep-tag II, Amp<sup>R</sup>, encodes AmiA Ct H67A | This work |
| pET-52b_*ct268*_H136A | Expression plasmid, T7 promotor, autoinducible in (DE3)-strains, N-terminal Strep-tag II, Amp<sup>R</sup>, encodes AmiA Ct H136A | This work |
| pET-52b_*ct268*_E205A | Expression plasmid, T7 promotor, autoinducible in (DE3)-strains, N-terminal Strep-tag II, Amp<sup>R</sup>, encodes AmiA Ct E205A | This work |

### Cloning

Expression plasmids pET-52b_*ct268,* pASK-IBA-3_*ct268* and pASK3C_*ct268* were constructed with In-Fusion® HD Cloning Kit (Takara Bio, Kusatsu, JPN) according to the manufacturer’s instructions using cloning primers listed in table 3. Mutagenesis was performed by using QuikChange Lightning Mutagenesis Kit (Agilent, Santa Clara, CA, USA). Mutagenesis primers are enlisted in table 3. For CRISPRi-constructs, *amiA*-targeting and non-targeting gRNA gBlock cassettes (Integrated DNA Technologies, Coralville, IA, USA) were inserted into BamHI-digested pBOMBL-Sa_dCas9vaa:L2 using the HiFi Assembly reaction master mix (NEB, Ipswich, MA, USA) according to the manufacturer’s instructions. gBlocks are listed in table 4. Chlamydial *amiA* with 6xH-tag was amplified by PCR with Phusion DNA polymerase (NEB, Ipswich, MA, USA) using *C. trachomatis* DNA as a template. PCR products were purified using PCR purification kit (Qiagen, Hilden, Germany). The product was inserted into SalI-digested pBOMBLCRia(*amiA*) by HiFi assembly reaction according to the manufacturer’s instructions. Plasmids were dephosphorylated with alkaline phosphatase (FastAP) (Thermo Fisher, Waltham, MA, USA) prior to HiFi reaction. Products of HiFi reactions were transformed into NEB-10ß cells and plated. Plasmids were isolated from colonies grown overnight in LB by using a plasmid mini-prep kit (Qiagen). Constructs were verified for correct size by digest, and inserts were sequenced.

**Table 3:**
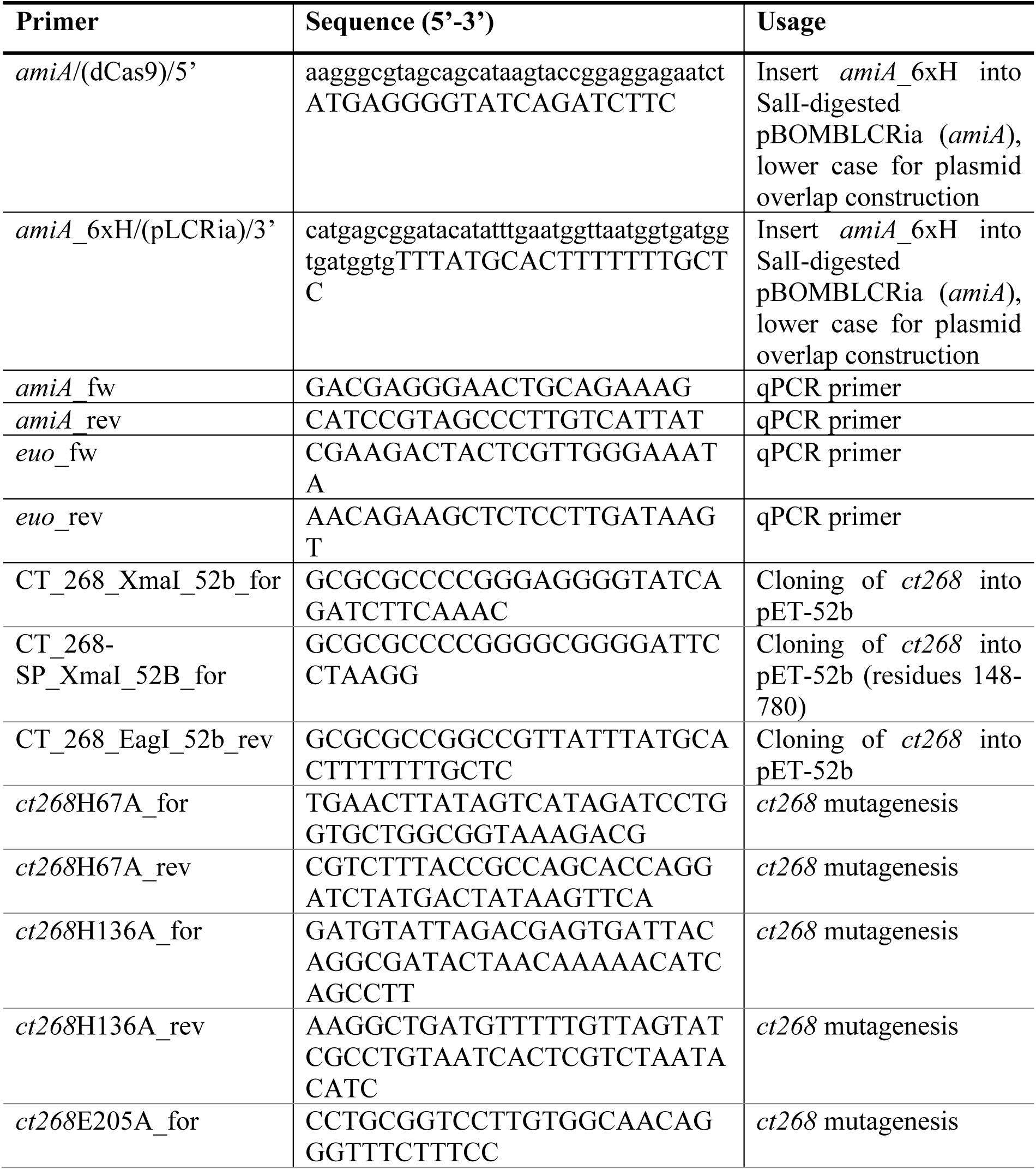
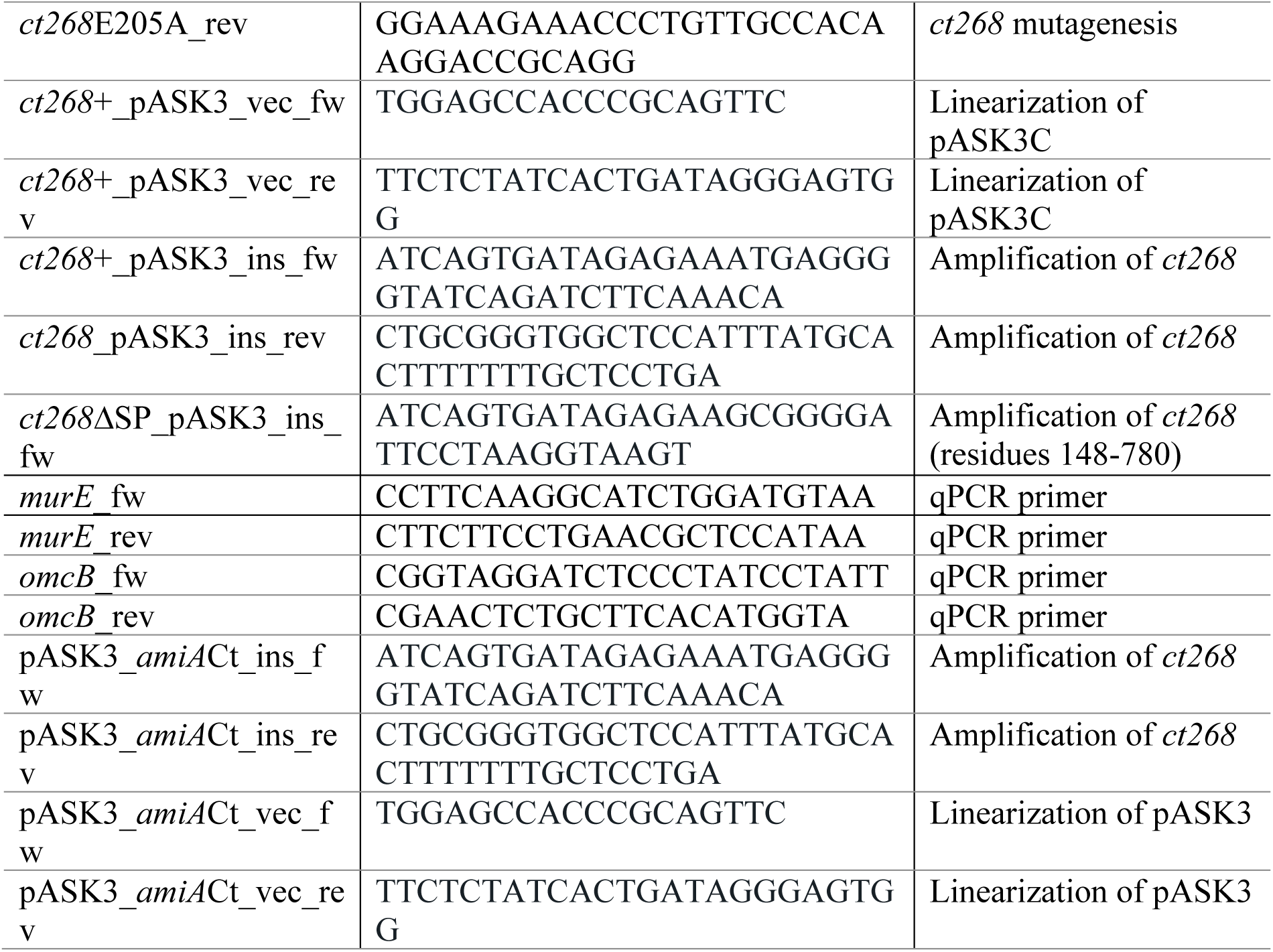
Primers.

**Table 4:**
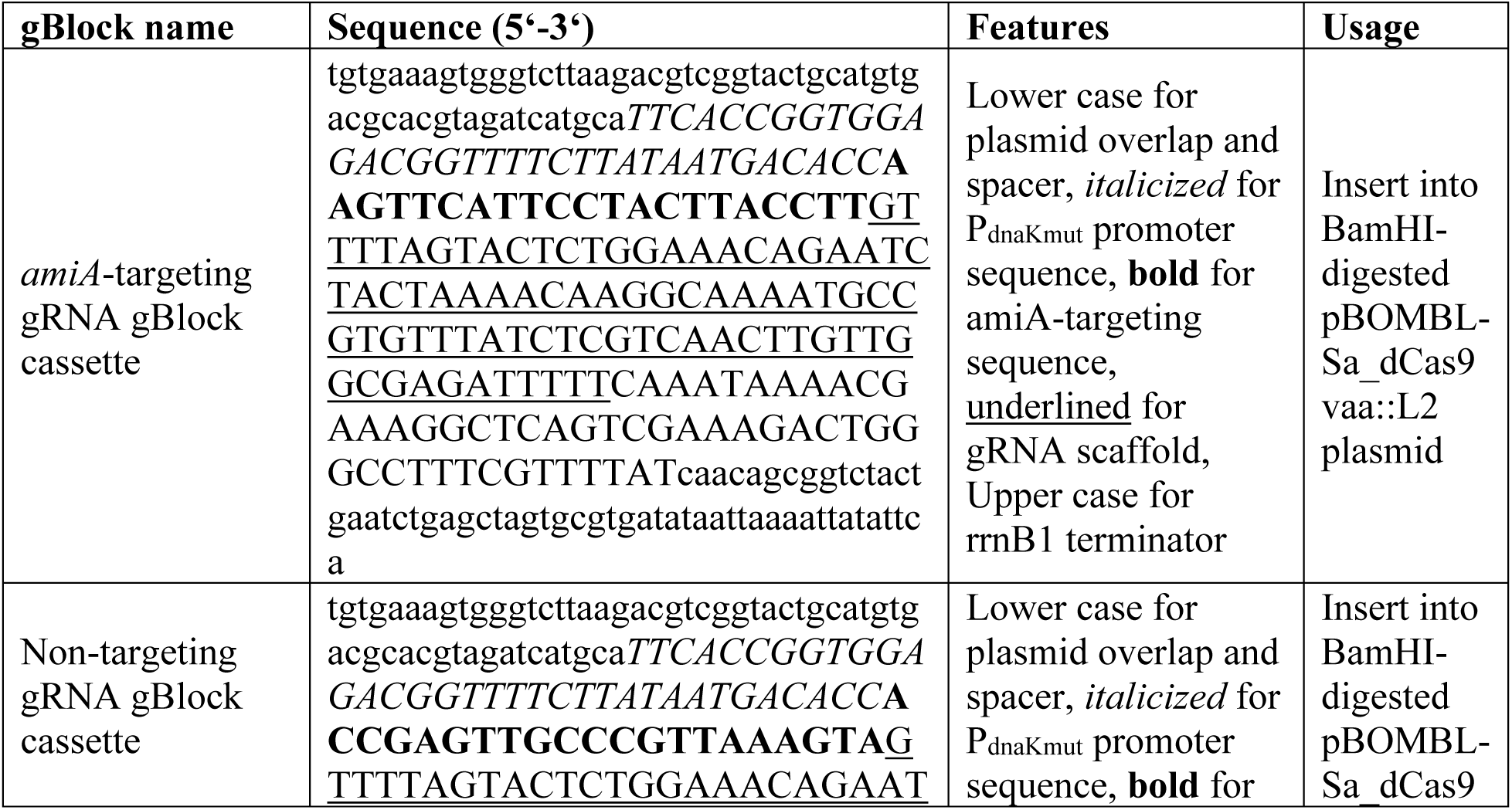
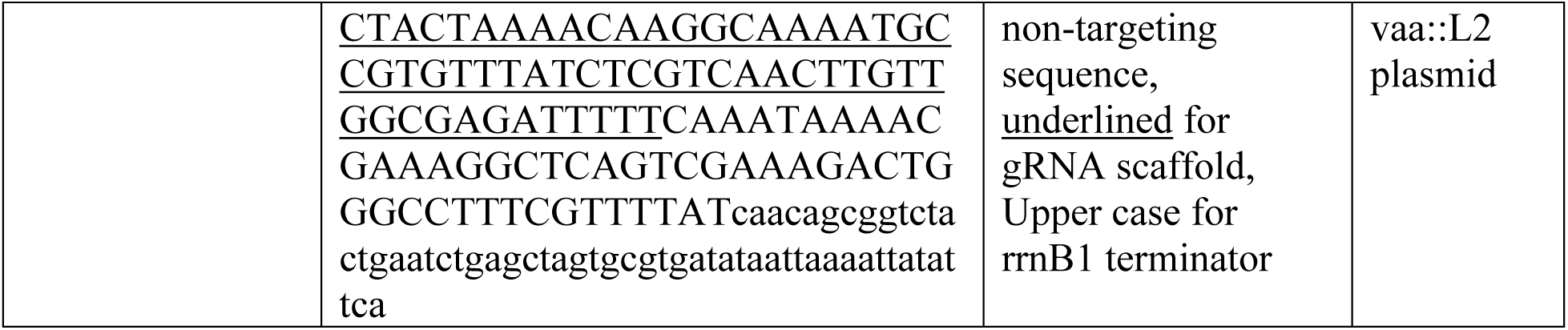
gBlocks.

### Protein purification

AmiA_Ct was overproduced in *E. coli* C43(DE3) cells during overnight incubation in TB-autoinduction medium [61] supplemented with 2g/L lactose, 0.5 g/L glucose, 10 mM MgSO_4_, 10 mM MgCl_2_ and 20 µM ZnCl_2_. Cells were resuspended in 50 mL buffer (25 mM MOPS pH 7.2, 1 M NaCl, 2 mM MgCl_2_) including 1 mM phenylmethyl sulfonyl fluoride (Thermo Fisher Scientific), 2 mg/mL polymyxin B (AppliChem, Darmstadt, Germany) and 20 U/mL benzonase A (Merck). Cells were disrupted by sonication, solubilized with 0.25 % CHAPS (Carl Roth, Karlsruhe, Germany) and incubated at 4 °C for 1 h before centrifugation at 135,000 x *g* for one hour at 4 °C. Purification was performed according to the manufacturer’s instructions for cleared lysates (IBA Lifesciences). Washing buffer was adjusted (25 mM MOPS pH 7.2, 500 mM NaCl, 2 mM MgCl_2_, 10 % (v/v) glycerol) and protein was eluted with 1 x BXT buffer buffer (IBA Lifesciences) containing 10 % (v/v) glycerol for cryoconservation. Dialysis was performed using D-Tube™ Dialyzer Maxi Tubes (Merck) in 1 L stirring dialysis buffer (25 mM MOPS pH 7.5, 300 mM NaCl, 2 mM MgCl_2_, 20 µM ZnCl_2_, 10 % Glycerol) at 4 °C overnight.

### Preparation of Remazol Brilliant Blue-stained peptidoglycan sacculi

Remazol Brilliant Blue-stained peptidoglycan sacculi (RBB-PGN) were used as a substrate in dye release assays and prepared as described in Farris *et al.* (2016) and Zhou *et al.* (1988) with modifications: *E. coli* W3110 was grown to an optical density of 0.6 in 4 L TB medium. Pelleted cells were resuspended in a final volume of 40 mL chilled H_2_O and boiled in 160 mL 5 % (w/v) SDS solution. Following overnight incubation at room temperature, sacculi were harvested by centrifugation at 9600 x *g* for 30 min. After washing with H_2_O, material was stained with 20 mM RBB solution in 0,25 M NaOH. Unbound dye was removed by multiple washing and centrifugation steps. RBB-PG was stored at –20 °C after lyophilization and dissolved in 0,2 % (v/v) Triton X-100 prior to use.

### Peptidoglycan binding assay

Pulldown assays were performed to determine binding affinity to PGN by AmiA_Ct. To eliminate non-functional and insoluble protein prior to the experiment, enzymes were pre-incubated for 30 min in buffer (50 mM Tris-HCl pH 8.5, 150 mM NaCl, 2 mM MgCl_2_, 20 µM ZnCl_2_). After centrifugation at 134,000 x *g* at 4 °C for 30 min, the protein-containing supernatant was added to to an equal volume of 2 mg/mL dissolved peptidoglycan of *B. subtilis* (Sigma-Aldrich, St. Louis, MO, USA). Tubes were incubated at 4 °C for 30 min and then centrifuged again. Samples taken from the supernantant represent unbound enzyme, while pellets were resuspended in 200 µL buffer and incubated on ice for 30 min, then centrifuged again. Supernatants and pellets were sampled and represent washing fraction and bound enzyme, respectively. Samples were analyzed by SDS-PAGE.

### Lysis Assay

Cell Lysis was monitored using a Tecan infinite M200 platereader (Tecan Group Ltd., Männedorf, Switzerland). *E. coli* JM83 was transformed with pASK3C-constructs encoding *amiA*_Ct and active site mutants **(Table 2)**. In a 96 well plate, cells were cultivated in LB-medium after inoculation with 1 % of overnight precultures. Gene expression was induced at OD 0.2-0.4 using 200 ng/mL AHT and then grown over night. Optical density measurements were performed every 15 min after brief shaking of the plate at constant temperature of 37 °C.

### Activity assays

*In vitro* activity assays using lipid II mDAP as a substrate were performed as described previously [31, 62]. Following overnight incubation, reaction products were analyzed by thin layer chromatography in Rick’s solvent (chloroform, methanol, H_2_O, ammonium hydroxide, 88:48:10:1) [63] and subsequent Hanessian’s stain [63]. Dye release assays were performed in a total volume of 50 µL containing 2 µM enzyme, 0,5 mg/mL RBB-PG, 50 mM Tris-HCl pH 8.5, 150 mM NaCl, 2 mM MgCl_2_, 20 µM ZnCl_2_ and 0,1 % (v/v) Triton X-100. For inhibitor assays, the DMSO-dissolved compounds were added in a 1:100 molar ratio (enzyme:inhibitor). pH was adjusted with MES pH 5.5, MES pH 6.5, MOPS pH 7.5 and CHES pH 9.5 for pH dependency analysis. Dye release assays were incubated at 37 °C for 1 h. Absorbance of the supernatant was measured at 595 nm.

### *In vivo* complementation assay

*In vivo* complementation assays with *E. coli* ADE24 harbouring the plasmid pASK-IBA3_*ct268* encoding AmiA_Ct including the native signal peptide were performed as described previously with modifications [31]. Cultures were incubated overnight and adjusted to an optical density of 0.5 before microscopy. Wild-type controls were supplemented with 0.5 % arabinose instead of glucose. A Zeiss AxioObserver Z.1 fluorescence microscope (Zeiss, Jena, Germany) with a 100 x oil immersion objective was used.

### Chlamydial transformation

2.6 x 10^6^ EBs of *C. trachomatis* L2 were incubated with 2 µg of plasmid in a total volume of 50 µL buffer (10 mM Tris, 50 mM CaCl_2_, pH 7.4) at room temperature for 30 min. Meanwhile, McCoy cell monolayers plated in 6 well plates were washed with 2 mL HBSS/well before 1 mL HBSS/well was added back. The transformation inoculum was mixed with 1 mL HBSS and added to cell monolayers for infection. For synchronous infection, cells were centrifuged at 400 *x g* for 15 min at RT and incubated at 37 °C for 30 min. Afterwards, inocula were aspirated and exchanged with DMEM + 10 % FBS + 10 µg/mL gentamycin. 8 hpi, media was replaced with DMEM containing the same amounts of FBS and gentamycin, and 1 µg/mL cycloheximide. 48 hpi, chlamydial transformants were harvested and used for infection of new McCoy monolayers until a stable population of transformants was established. Then, EBs were harvested and stored in sucrose-phosphate buffer (2SP) at – 80 °C. Transformants were verified by isolation of plasmid DNA, restriction digest and sequencing.

### Determination of genetic effects of CRISPRi-mediated knockdown of *amiA* by RT-qPCR

HeLa-2 cells were plated in 6 well plates in triplicate wells per test condition and were infected with *C. trachomatis* L2 transformants at a MOI of 0.1. At 10 hpi, construct expression was induced or not with 5 nM aTc. To collect RNA, infected cell monolayers were harvested with 500 µL TRIzol™ (Invitrogen™, Carlsbad, MA, USA). To digest DNA, samples were treated with Turbo DNAse from Turbo DNA-free™ kit (Invitrogen™). During RT-PCR, cDNA was transcribed from RNA using SuperScript™ III Reverse Transcriptase (Invitrogen™) according to the manufacturer’s instructions. cDNA was diluted 10 x with nuclease-free water and frozen at – 80 °C. Equal volumes of of cDNA were used in a total volume of 25 µL for qPCR with SYBR green master mix (Applied Biosystems) and quantified on a QuantStudio 3 (Applied Biosystems/Thermo Fisher) using standard amplification cycle with melting curve analysis. Results were compared to a standard curve generated against purified *C. trachomatis* L2 genomic DNA. DNA samples were collected and purified using the DNeasy blood & tissue kit (Qiagen) according to the manufacturer’s instructions. Equal DNA quantities were used in qPCR with an *euo* primer set to quantify chlamydial genomes to normalize respective transcript data. qPCR primer sequences are listed in **Table 3**. Results were normalized for efficiency, and Student’s t-test was used to compare gene expression levels of uninduced and induced samples at 10 hpi, 14 hpi and 24 hpi. To verify construct expression, infected cells plated on coverslips were fixated with 100 % methanol for 5 min. For staining, a primary goat antibody directed against chlamydial major outer membrane protein (MOMP) (Meridian, Memphis, TN), a donkey anti-goat antibody conjugated to Alexa Fluor 488 (Invitrogen™), a primary rabbit antibody directed against dCas9 (abcam, Cambridge, UK, ab203933) and a donkey antirabbit antibody conjugated to Alexa Fluor 594 (Jackson ImmunoResearch, Ely, UK: 711-586-152) were used. Images were acquired at 100 x magnification on a Zeiss AxioImager.Z2 (Zeiss) microscope with an Apotome2 and an Axiocam 506 6 MP digital monochrome camera.

### Inclusion-forming-unit assays to determine effects of *amiA* knockdown on infectious progeny

Infectious progeny was determined by reinfection of a cell monolayer with chlamydial EBs harvested from a primary infection. HeLa cells were infected with chlamydial transformants bearing constructs of interest. After induction or not with 5 nM aTc 10 hpi, EBs were harvested at 24 hpi by scraping cells in 2SP buffer. Samples were lysed during one freeze-thaw cycle, serially diluted and used to infect a new HeLa monolayer. To verify construct expression, infected cells plated on coverslips were fixated with 100 % MeOH for 5min and immunostained for MOMP and dCas9 and imaged as described previously. The secondary infection was maintained for 24h, then fixated with 100 % MeOH for 5 min and immunostained against MOMP. IFU titers were obtained by calculating the total number of inclusions per field of view based on counts of 15 fields of view.

### Chlamydial peptidoglycan labelling, imaging, and quantification

EDA-DA incorporation assays were performed as previously described [12]. Briefly, EBs from *C. trachomatis* transformants were used to infect cell monolayers (previously seeded on 24-well coverslips the day before) via rocking incubation for 2 hours at 37 °C under tissue culture conditions. EDA-DA was then added, at a concentration of 4mM, at either 2 hpi or 23 hpi. At the assayed designated endpoint infected cells were then fixed with 100% Methanol at room temperature (RT) for 5 minutes, washed x3 with PBS, further permeabilized with 0.5% Triton X for 5 minutes, washed an additional x3 with PBS, and blocked with 3% BSA for 1 hour at RT. Subsequently, cell coverslips were incubated with 100ul of Click-iT^TM^ Cell Reaction Buffer (ThermoFisher) reaction mix, according to the manufacturer’s instructions. 100ul of reaction mix was used per coverslip with 1ul of Alexa Fluor^TM^ 488 azide (Invitrogen). Bacteria were counter labelled with chlamydial anti-MOMP (LS-C79219) at 1:500 and secondary anti-Goat Alexa Fluor^TM^ 594 (Invitrogen) at 1:1000. Coverslips were mounted on slides with Prolong Gold Antifade Mounting Media and stored at 4 °C prior to imaging. Slides were imaged on a Zeiss ELYRA PS.1 in SIM mode. zStacks were collected and representative images presented as 3D rendered composites in Figure 5. EDA-DA-labelled object quantification was performed as previously described [24]. ImageJ (FiJi) software addon “3D Objects Counter” was used to delineate individual PG-containing objects and assess their relative sizes and fluorescent intensities. Data presented represent all EDA-DA-labeled objects present in zStacks of 15 inclusions per condition / strain examined spanning 2 biological replicates. Object size thresholding was restricted to objects larger than 0.001 μm^3^ and smaller than 0.15 μm^3^.

## Acknowledgments

This work was supported in part by funding from the National Institutes of Health (NIGMS) to SPO (1R35GM151971) and GWL (1R35GM138202). Funding was also provided by the Deutsche Forschungsgemeinschaft (DFG, German Research Foundation), project-ID 398967434 —TRR261 (BH) and 390536577 (BH), and the funding scheme FEMHABIL, Medical Faculty, University of Bonn (BH). The funders had no role in study design, data collection and interpretation, or the decision to submit the work for publication. The opinions and assertions expressed herein are those of the author(s) and do not necessarily reflect the official policy or position of the Uniformed Services University or the Department of Defense.

## Supplemental Information

**Suppl. figure 1:**
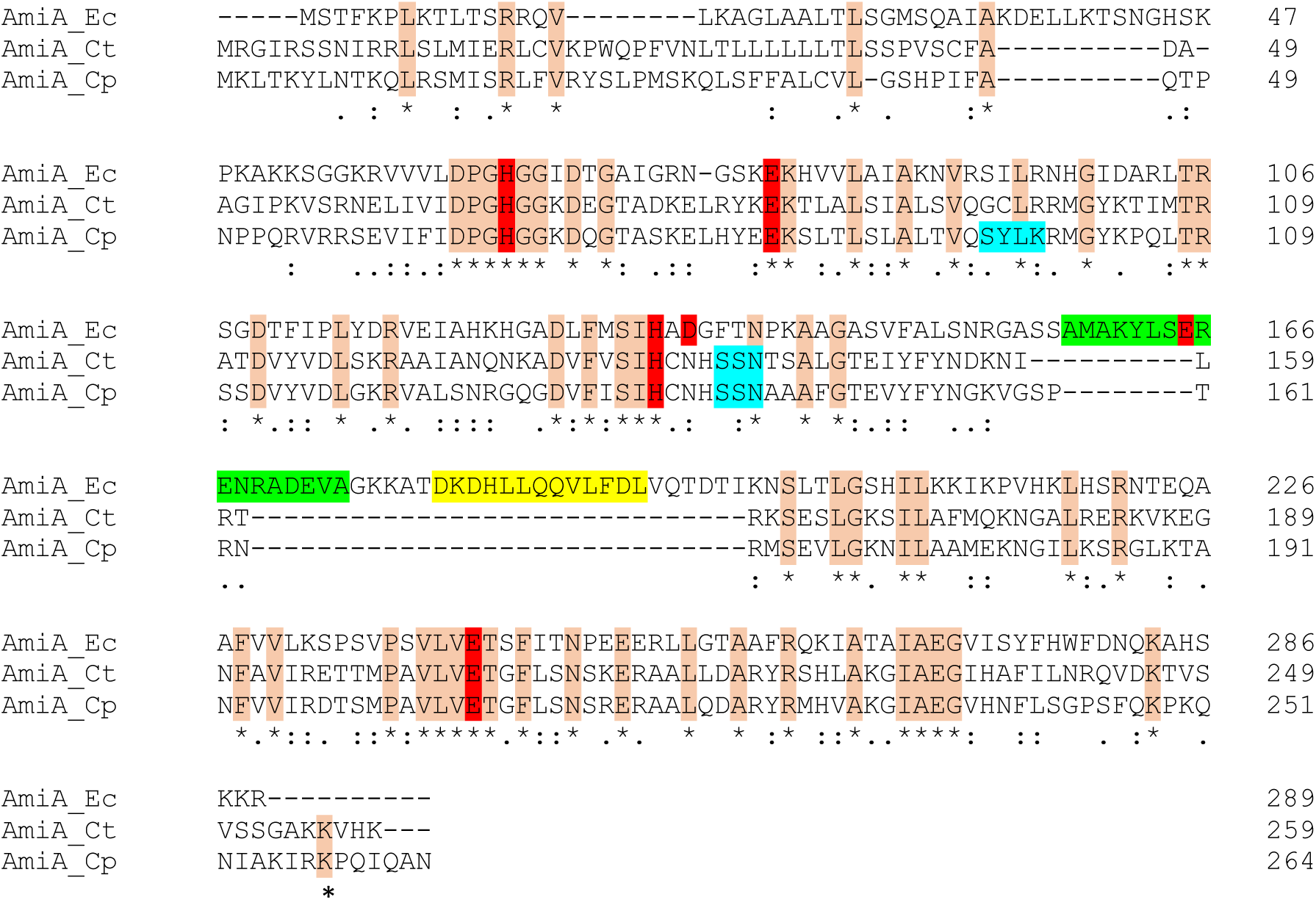
Amino acid sequence alignment of AmiA homologs of *C. trachomatis, C. pneumoniae* and *E. coli*. Red: active site residues. Orange: Identical residues. Blue: PBP motifs. Green: blocking helix. Yellow: interaction helix. Below the alignment, asterisks (*) indicate fully conserved residues between input sequences. Colons (:) indicate conservation of strongly similar properties, while periods (.) indicate conservation of weakly similar properties based on Gonnet PAM 250 matrix used in Clustal Omega alignments.

**Suppl. figure 2:**
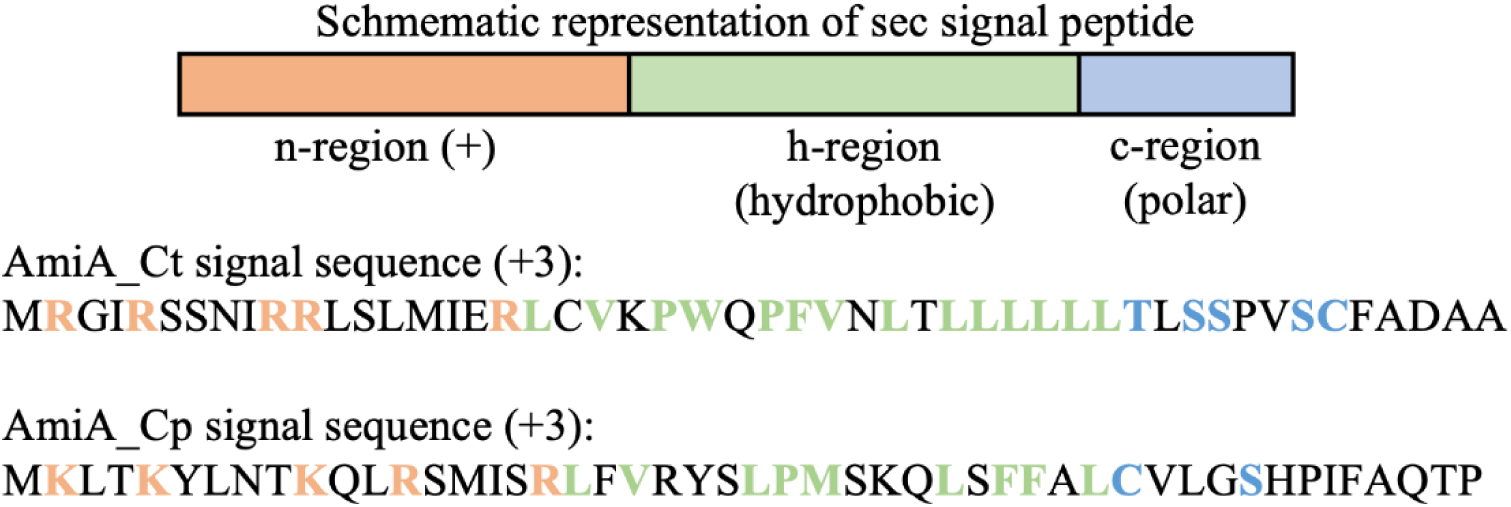
Schematic representation of sec-system signal peptides and signal peptide sequences of AmiA_Ct and AmiA_Cp. Positively charged (+) amino acids in the n-region are marked in red, while hydrophobic amino acids of the h-region are shown in yellow. Polar amino acids of the c-region are depicted in blue. The consensus motif for recognition of the cleavage site by signal recognition particle (SRP) and signal peptidase is A-x-A. Matching amino acids are shown in grey. This suggests a cleavage between residues 47 and 48 in AmiA_Ct, which was also predicted by SignalP 5.0.

